# Bacterial filamentation drives colony chirality

**DOI:** 10.1101/2021.03.11.434908

**Authors:** Andrés Aranda-Díaz, Cecilia Rodrigues, Alexandra Grote, Jiawei Sun, Carl Schreck, Oskar Hallatschek, Anton Souslov, Wolfram Möbius, Kerwyn Casey Huang

## Abstract

Chirality is ubiquitous in nature, with consequences at the cellular and tissue scales. As *Escherichia coli* colonies expand radially, an orthogonal component of growth creates a pinwheel-like pattern that can be revealed by fluorescent markers. To elucidate the mechanistic basis of this colony chirality, we investigated its link to left-handed, single-cell twisting during *E. coli* elongation. While chemical and genetic manipulation of cell width altered single-cell twisting handedness, colonies ceased to be chiral rather than switching handedness, and anaerobic growth altered colony chirality without affecting single-cell twisting. Chiral angle increased with increasing temperature even when growth rate decreased. Unifying these findings, we discovered that colony chirality was associated with the propensity for cell filamentation. Inhibition of cell division accentuated chirality under aerobic growth and generated chirality under anaerobic growth. Thus, regulation of cell division is intrinsically coupled to colony chirality, providing a mechanism for tuning macroscale spatial patterning.

## Introduction

An object is chiral if it is distinguishable from its mirror image. Chirality is prevalent throughout nature at all scales, and stereoisomers are often functionally distinct, from our left and right hands to L- and D-amino acids that are used for metabolism/translation and bacterial cell-wall synthesis, respectively. Chirality is manifest in polymers that form helices, such as bacterial flagella and cytoskeletal filaments (1, 2). Chirality can also be an intrinsic property of individual cells; for instance, myosin in *Drosophila* can reverse handedness in cells, which feeds forward to affect organ handedness (3). Chirality drives the development of left-right asymmetry generation in organs of *Drosophila melanogaster* (4) and in the *Caenorhabditis elegans* embryo (5). Plants twist as they grow, and mutants in SPIRAL2 change that twist from left-to right-handed; this handedness reversal is coupled to a switch from anisotropic growth to isotropic growth (6). However, it is largely unknown how chirality at the tissue and organismal scales is linked to cellular and molecular properties. Here, we investigate the link between chirality and growth at the micron scale of individual bacterial cells and chirality at the millimeter scale of colonies, visible to the naked eye.

Many rod-shaped bacterial cells exhibit twisting at the single-cell level during growth. In the Gram-negative bacterium *Escherichia coli*, growth occurs along the body of the cell and not at the poles (7). As the two ends move apart from one another, they also rotate in opposite directions, representing left-handed twist (Fig. 1A) (8). Twisting has also been observed in the Gram-positive *Bacillus subtilis* (9), in a right-handed manner (8). Bacterial growth requires expansion of the cell wall, a rigid macromolecule composed of crosslinked glycan strands (10) that is necessary and sufficient for cell-shape determination (11). A biophysical model of cell-wall growth quantitatively predicted the degree of cell twisting generated by a helical pattern of insertion, which produced cell-wall material with the opposite handedness (8, 12). The bacterial actin homolog MreB (13), which is responsible for the spatiotemporal patterning of cell wall material (7), was required for cell twisting (8). The small molecule A22 depolymerizes MreB; at high concentrations, cells eventually round up and lyse (14). Twisting of *E. coli* cells is tunable: as cells widen under increasing sublethal levels of A22 treatment (15), the angle of MreB motion, which is thought to signify the placement of new strands of cell wall material (16), rotates and ultimately adopts an angle on the opposite side of the line perpendicular to the long axis of the cell, signifying a gradual conversion of twisting from left-to right-handed (15).

**Figure 1:**
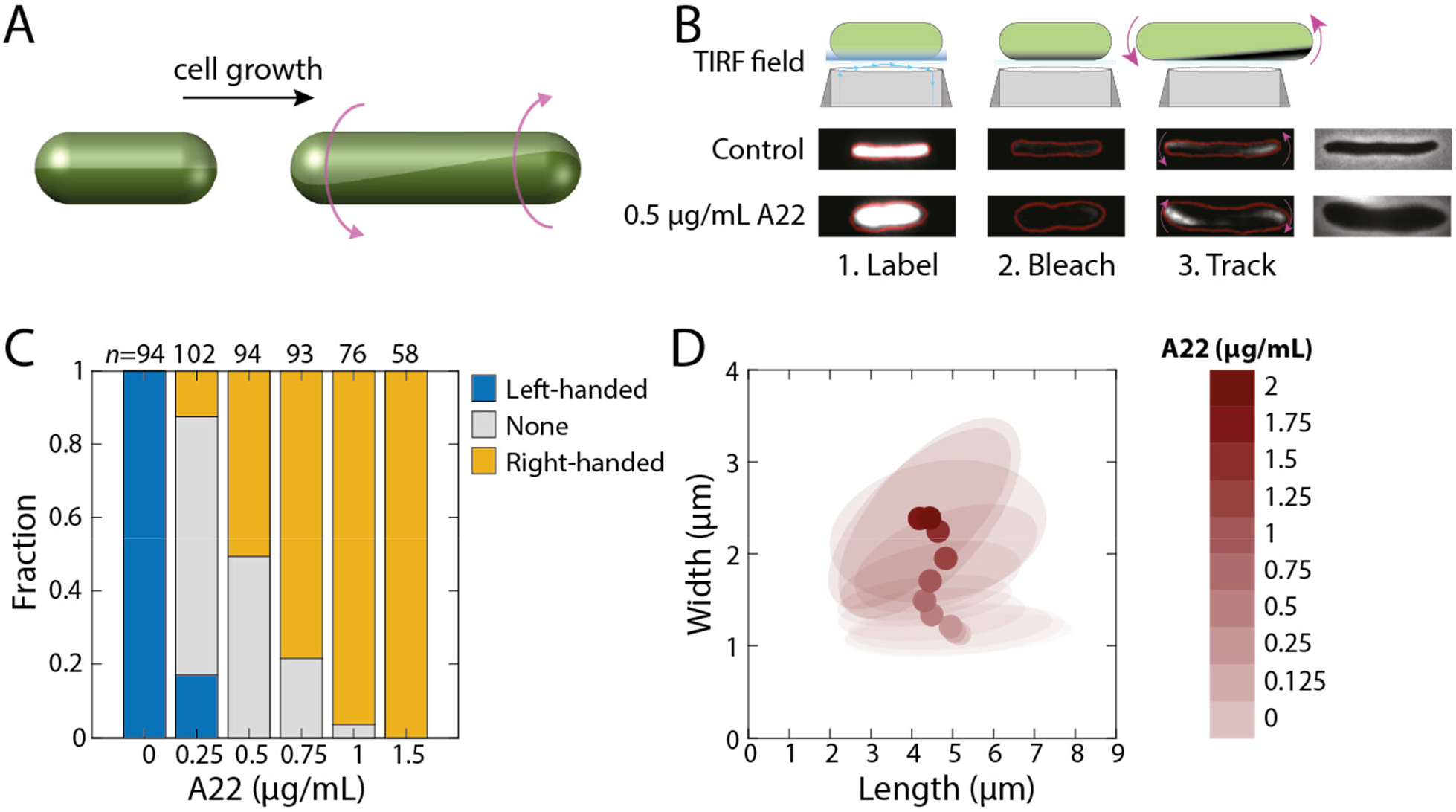
A22 reverses twisting handedness and alters cellular dimensions in *E. coli* DH5α. A) Schematic of left-handed twisting during elongation of a rod-shaped cell. B) In the Twist-n-TIRF method, the cell surface is labeled and the side of the cell closest to the cover slip is bleached using a TIRF microscope. During subsequent TIRF imaging at lower intensity, unbleached parts of the cell appear associated with cell twisting (Methods). Rightmost panel: phase-contrast images. A22-treated *E. coli* DH5α cells treated with A22 are typically shorter and wider than untreated cells and twist with opposite handedness during the tracking step; note the appearance of fluorescence on the lower left and upper right in the A22-treated cell, as opposed to the upper left and lower right in the untreated cell. C) In the absence of A22, virtually all *E. coli* DH5α cells exhibit left-handed twisting, while cells treated with 1.5 µg/mL A22 exhibit right-handed twisting. The number of cells (*n*) is indicated above each bar. D) *E. coli* DH5α cell width during log-phase growth in liquid increases as a function of A22 concentration. For concentrations <1 µg/mL, cell length decreases with increasing A22 concentration. Circles represent mean dimensions and ellipses represent the covariance matrix of length and width. *n*>50 cells were quantified for each condition.

Bacterial colonies can also exhibit chirality during growth (17-19). This effect is striking in experiments investigating range expansions, in which otherwise genotypically and phenotypically identical cells producing fluorescent proteins of two different colors (purely for the purposes of distinguishing genotypes) are inoculated onto an agar plate to grow into a colony. As the colony expands, cells on the exterior have preferential access to uncolonized surface area and nutrients, leading to spatial segregation of the two fluorophores in well-defined “sectors” that expand outward, ultimately producing a pinwheel pattern. Boundaries between these sectors provide a frozen record of colony growth that displays a chiral angle (19). On top of the wiggling motion of boundaries between sectors, the boundaries of many species such as *E. coli* exhibit deterministic chirality in the form of expansion biased along the edge of the colony (17). Ultimately, cells themselves are likely to generate the observed behavior at the colony scale, although colony chirality must also be dependent on environmental conditions such as adhesion and surface wetness, both of which dictate cell motility. Indeed, *E. coli* colony chirality was shown to be mediated by close interactions between cells and the surface, with expression of pili and other adhesive structures suppressing chirality, and agar stiffness affecting chirality (20). There are also strain-specific differences in colony chirality (20), even though twisting at the single-cell level is relatively constant (8). Previous studies that attempted to explore the relationship between cell shape or cell-wall synthesis and colony chirality made comparisons between different species (20). However, a systematic interrogation of the links between single-cell properties and colony chirality through environmental, genetic, and physiological perturbations has not been undertaken. In particular, it is critical to employ a strategy that tunes behaviors such as twisting in a single organism in order to probe potential couplings with colony chirality.

Here, we set out to determine the relationship between cell shape, twisting and handedness at the single-cell level, and macroscopic colony chirality. Using single-cell and colony imaging, we found that A22 treatment and anaerobic growth inhibited growth and reduced colony chirality to near zero, making it unclear whether single-cell twisting was responsible for the change in chiral angle. Cells at the edge of the colony adapted to A22 treatment by reducing their width and length. Chiral angle increased with increasing temperature, even at high temperatures that caused a decrease in growth, indicating that growth rate does not determine colony chirality. Across all conditions, the presence of chiral colonies was associated with filamentous cells at the edge of the colony, and antibiotic inhibition of division resulted in enhanced chirality. These results reveal a complex connection between single-cell dimensions and population level spatial patterns, underscoring the role of cell division in determining colony chirality.

## Results

### *E. coli* DH5*α* cells exhibit similar single-cell twisting as MG1655

Chirality in *E. coli* colonies is readily observed when using strain DH5*α* (17, 19), with a clockwise rotation when viewed from the top. A recent comparison between *E. coli* strains revealed that on low-salt LB agar, *E. coli* MG1655 colonies also rotate clockwise, but exhibit less pronounced chirality than DH5*α* colonies (20). To determine whether this difference in chirality could be due to differences in twisting at the single-cell level (Fig. 1A), we utilized our previously developed Twist-n-TIRF method (15). In this method, cells are treated with the beta-lactam antibiotic cephalexin to block cell division, allowing for easier visualization of twisting. The cell wall is labeled uniformly with a stationary dye (in this case, Wheat Germ Agglutinin (WGA) labeled with Alexa Fluor 488 (7)), and then the bottom of the cell is bleached by a TIRF laser. As the cell subsequently grows, twisting causes bright, unbleached regions to rotate into the TIRF imaging plane (Fig. 1B), and the handedness and degree of twisting can be computed from the direction and rate of fluorescence appearance, respectively (15). Virtually all DH5*α* cells clearly twisted in a left-handed manner on LB agarose pads (Fig. 1C), similar to MG1655 on EZ-RDM pads (15) and consistent with our previous study employing beads bound to the outer membrane (8). Thus, left-handed twisting is conserved across *E. coli* strains and growth media.

Given the central role of MreB in single-cell twisting (8, 15), we next sought to quantify the effects of A22 treatment on DH5*α*. We treated log-phase DH5*α* cells expressing YFP or CFP from a plasmid for 3 h with a range of A22 concentrations from below to above the minimum inhibitory concentration (∼1 µg/mL) in LB (Methods). At higher A22 concentrations, Twist-n-TIRF measurements revealed an increasing fraction of DH5*α* cells that exhibited right-handed or ambiguous twisting (Fig. 1C), similar to the handedness reversal of MG1655 (15). Cell shape also changed as expected from previous experiments with MG1655 (15): DH5*α* cells increased in width and decreased in length as A22 concentration was increased (Fig. 1D). Thus, DH5*α* exhibits similar changes in twisting and cell shape across A22 concentrations as MG1655.

### A22 treatment reduces colony chirality

The inversion of twisting handedness due to A22 treatment provides a qualitative change in cellular behavior through which to interrogate the connection between single-cell and colony-level behaviors. If single-cell twisting is the determinant of colony chirality, we should see a change from clockwise to counter-clockwise rotation within colonies as single-cell twisting changes with increasing A22 concentration. To test this hypothesis, we grew mixed colonies of YFP- and CFP-expressing DH5*α* on LB plates at 37 °C with a range of A22 concentrations and imaged the colonies after 7 days of growth. As expected, the frozen record revealed that colonies quickly developed into sectors of single colors (Fig. 2A). We segmented the images, identified sector boundaries, plotted the change in angle against a function of colony radius, and computed the chiral angle (Fig. 2A, Methods). In the absence of A22, the chiral angle was *θ*=6.4° (Fig. 2B,C), similar to previous measurements (17, 20). A22 treatment reduced the chiral angle to approximately zero (Fig. 2B,C), but even at high concentrations there was not clear evidence for reversal of handedness, in seeming contradiction to our hypothesis.

**Figure 2:**
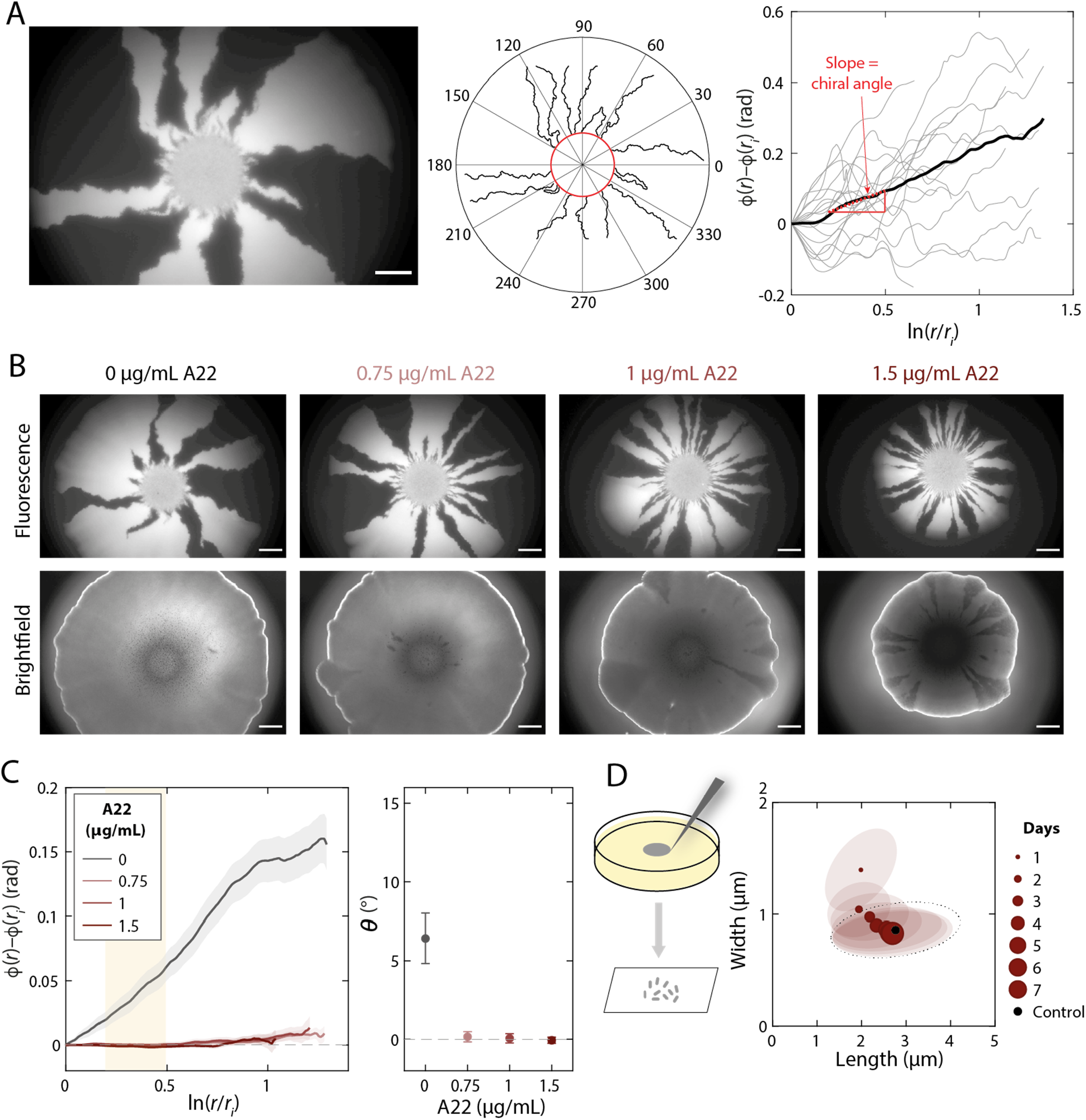
A22 treatment reduces colony chiral angle. A) Genetic demixing during *E. coli* DH5*α* colony growth results in monoclonal sectors (left). Although the shape of the boundaries between light (YFP) and dark sectors (middle) appeared stochastic, quantitative image analysis revealed an overall clockwise rotation of sector boundaries when viewed from the top (air interface). Right: the slope (red) of the mean (thick black curve) is defined as the chiral angle. Scale bar: 1 mm. B) Images of typical colonies after 7 days of growth on plates with various concentrations of A22 illustrate that colony growth is hindered by A22. The bright outline in the brightfield images denotes the colony border. Sector boundaries were straighter at higher A22 concentrations. Scale bars: 1 mm. C) Left: mean rotation of sector boundaries at various A22 concentration. Right: the chiral angle decreases at higher A22 concentrations. Each data point is the average of *n*≥5 colonies. Error bars represent 1 standard deviation. D) During colony growth in the presence of 1.5 µg/mL A22, cellular dimensions gradually revert back to those of cells grown in colonies in the absence of A22, suggesting adaptation to A22.

In a recent study, we showed that *E. coli* adapts to the cell widening effects of A22 treatment by increasing expression of *mreB*, resulting in a subsequent decrease in cell width (21). If adaptation that altered cellular dimensions occurred during colony growth, such changes could have confounded our above test of the effect of A22 and single-cell twisting on colony chirality. To test whether cells on LB plates with A22 changed morphology over time, we sampled cells from the edge of colonies once per day and imaged the cells to quantify their dimensions (Methods). As we suspected, cell width decreased steadily over the course of a week, reaching a relatively constant width displayed by cells on plates without A22 (Fig. 2D); coincident with the width decrease, length gradually increased (Fig. 2D). For the widest cells (∼1.4 µm), the cell width measured on day 1 on LB+1.5 µg/mL A22 was equivalent to the cell width in liquid LB+0.5 µg/mL A22 (Fig. 1D). Under this condition, ∼60% of cells exhibited no or ambiguous twisting (Fig. 1C), consistent with the lack of chirality at the colony level. Thus, adaptation of cell shape on plates prevents a direct test of whether A22-mediated reversal of single-cell twisting handedness necessarily reverses colony-chirality handedness.

### Heterologous expression of a foreign cell-wall synthesis enzyme reverses single-cell twisting but colonies do not exhibit chirality

To circumvent adaptation associated with A22 treatment, we sought an alternate mechanism for altering cell width and single-cell twisting. We previously showed that heterologous expression of *mrdA*, which encodes the essential transpeptidase PBP2 (22), from *Vibrio cholerae* caused *E. coli* MG1655 cells lacking the endogenous *mrdA* gene to widen and to reverse twisting handedness similar to A22 treatment (15). We constructed DH5*α* Δ*mrdA* strains with constitutive expression of *V. cholerae mrdA* (Vc-*mrdA*) and a plasmid coding for CFP or YFP from a parental strain DH5*α*-E, resulting in DH5*α*-E Δ*mrdA* Vc-*mrdA* CFP and DH5*α*-E Δ*mrdA* Vc-*mrdA* YFP (Methods). The mean cell width of these strains was significantly larger than DH5*α*-E or the DH5*α*-H strain used in the experiments above (Fig. 3A,B). DH5*α*-E Δ*mrdA* Vc-*mrdA* cells were highly sensitive to cephalexin, resulting in rapid lysis during Twist-n-TIRF experiments that precluded measurement of single-cell twisting. Unlike A22-treated wild-type cells, untreated fluorescent DH5*α*-E Δ*mrdA* Vc-*mrdA* cells remained wider and shorter than wild-type cells over 5 days of growth in colonies (Fig. S2). Colonies displayed highly reduced chiral angle (Fig. 3C,D), leaving it unclear whether reversal of handedness at the single-cell level results in reversed handedness of colony chirality.

**Figure 3:**
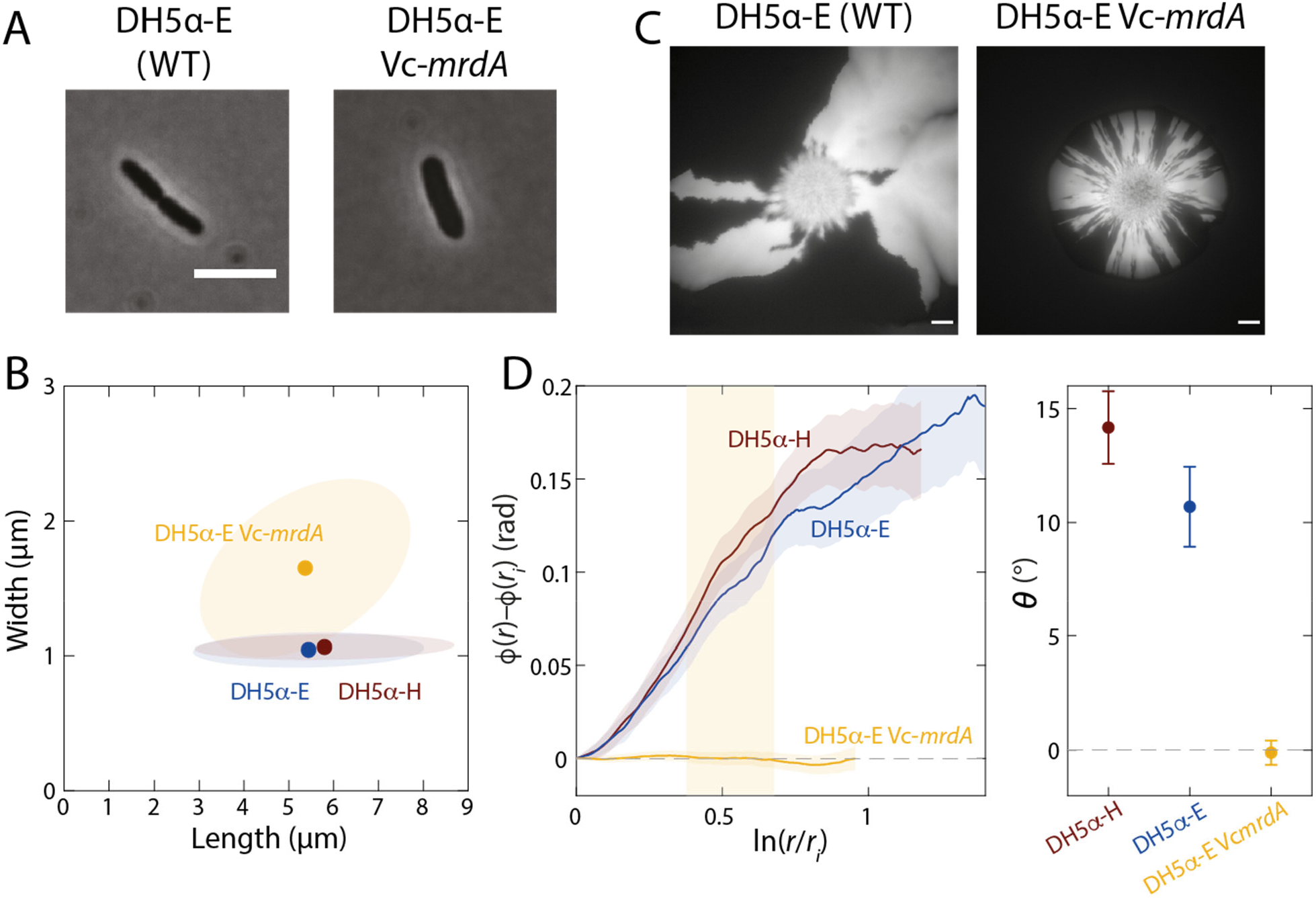
A genetic perturbation that causes cells to become wider eliminates colony chirality. A,B) Heterologous expression of the *mrdA* gene from *Vibrio cholerae* (Vc-*mrdA*) in the *E. coli* DH5*α*-E Δ*mrdA* background increases cell width during log phase in liquid growth relative to wild-type DH5*α*-H or DH5*α*-E cells. In (B), circles represent mean values and ellipses represent the covariance of width and length. *n*>50 cells were quantified for each strain. Scale bar: 5 µm. C,D) Colony chirality is reduced by heterologous expression of Vc-*mrdA*. (C) Image of typical colonies after 7 days of growth. Scale bars: 1 mm. (D) Left: mean rotation of sector boundaries for each strain. Right: the chiral angle is essentially zero for the Vc-*mrdA* strain. Each data point is the average of *n*≥5 colonies. Error bars represent 1 standard deviation.

### Quenching of colony chirality between two surfaces is likely due to lack of twisting in anaerobic environments

We hypothesized that colonies sandwiched between two agar surfaces should not exhibit any chirality based on symmetry considerations, independent of changes in cell shape. To test this hypothesis, we inoculated a droplet of CFP-expressing and YFP-expressing DH5*α* cells as before, allowed the droplet to dry, and then placed another large agar pad on top of the agar plate (Fig. 4A). The colony continued to expand between these two surfaces, presumably due to the lack of agar crosslinking between the two pads as compared to within the pads. Cells emitted very little fluorescence, which we surmised was due to their lack of oxygen; anaerobic conditions prevent maturation of fluorescent proteins (23). GFP matures more readily at low temperatures (24), hence we incubated sandwiched colonies at 4 °C (Methods) and then quantified the fluorescence patterns. The sector boundaries were essentially achiral (Fig. 4A,B, S3).

**Figure 4:**
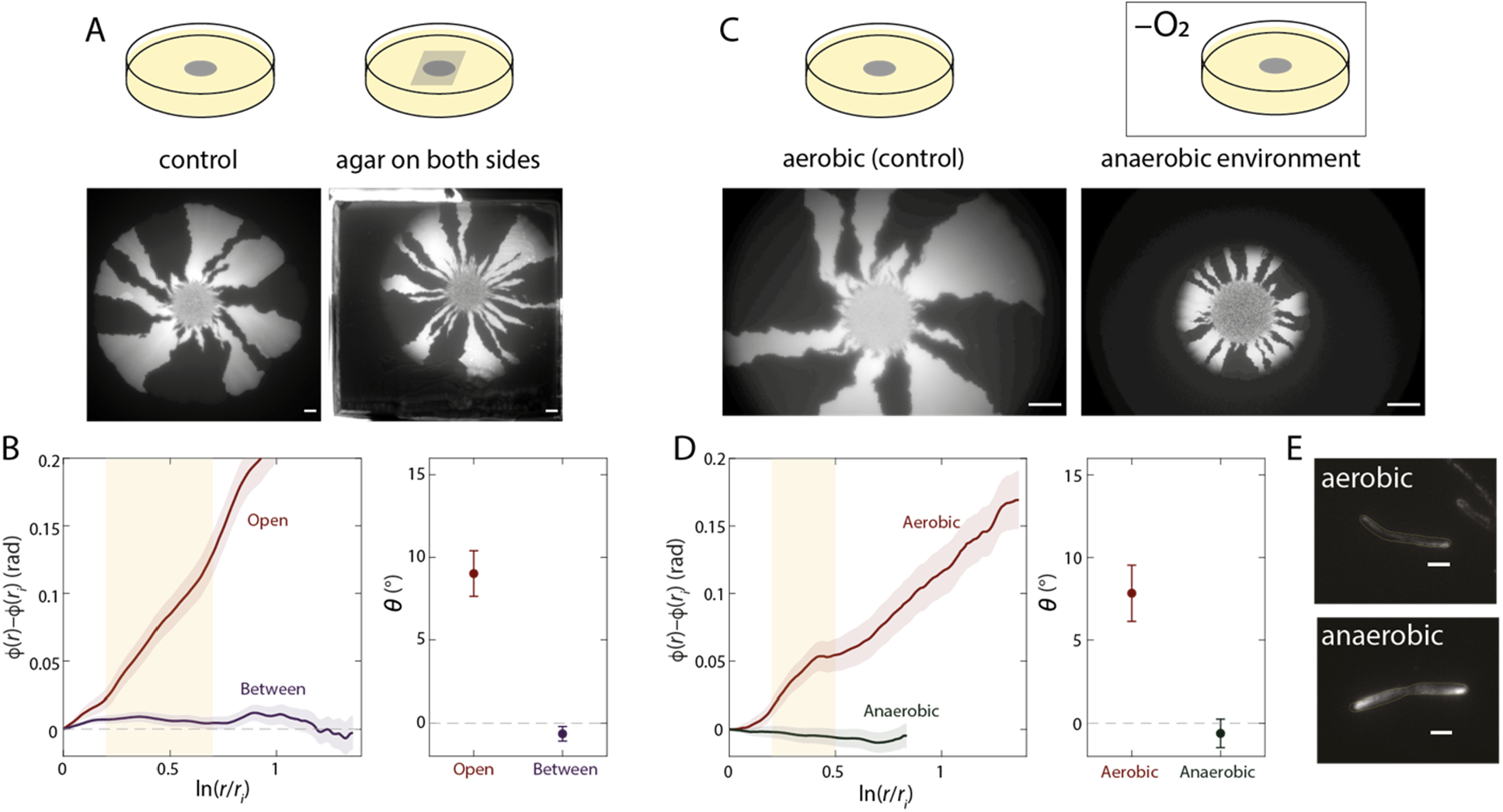
Colony chirality is decreased during growth when sandwiched between agar surfaces or in anaerobic conditions. A) Top: schematic of control experiments with an air-agar interface (“Open”, left) and sandwiched between two agar surfaces (“Between”, right) (Methods). Bottom: representative colonies for each condition. Sector boundaries were straighter in the sandwiched colony. Scale bars: 1 mm. B) Left: mean rotation of sector boundaries in each condition in (A). Right: the chiral angle is essentially zero for sandwiched colonies. Each data point is the average of *n*≥5 colonies. Error bars represent 1 standard deviation. C) Top: schematic of growth in aerobic conditions and in an anaerobic chamber. Bottom: representative colonies for each condition. Sector boundaries were straighter during anaerobic growth. Scale bars: 1 mm. D) Left: mean rotation of sector boundaries in each condition in (C). Right: the chiral angle is essentially zero during anaerobic growth. Each data point is the average of *n*≥5 colonies. Error bars represent 1 standard deviation. E) DH5*α* cells still exhibit twisting at the single-cell level during anaerobic growth at 37 °C, as revealed by Twist-n-TIRF. For both aerobic and anaerobic growth, all cells whose handedness could be reliably classified were left-handed (*n*=43, aerobic; *n*=31, anaerobic). Scale bars: 2 µm.

To test whether the absence of chirality was due to the restoration of symmetry at the interface or to the depletion of oxygen, we grew a mixed colony on a single agar surface in anaerobic conditions. We observed little to no chirality (Fig. 4C,D), indicating that anaerobic growth is sufficient to abolish colony chirality. Colonies were smaller after 7 days of anaerobic growth compared with aerobic growth. To test whether anaerobic growth abolished single-cell twisting, we performed Twist-n-TIRF in anaerobic conditions (Methods). Cells continued to twist in a left-handed fashion (Fig. 4E), thus lack of colony chirality in anaerobic conditions is not due to lack of single-cell twisting. Instead, these findings suggest that another, oxygen-dependent factor influenced the level of colony chirality.

### Temperature alters growth rate and chirality without changing cell twisting

After 7 days, colonies grown on LB+A22 (Fig. 2A) or anaerobically (Fig. 4C) were significantly smaller than when grown aerobically on LB alone (Fig. 2A, 4C). Thus, we sought to test whether growth rate was a major determinant of colony chirality. Temperature is well known to modulate growth rate (25). In liquid, the maximum growth rate of DH5*α* increased as a function of temperature up to an optimal temperature of ∼42 °C (Fig. 5A, S4). We grew mixed colonies across a range of temperatures and imaged them after 7 days (Fig. 5B). Colony size was also temperature-dependent, with smaller colonies at 30 and 42 °C than at 37 °C (Fig. 5A). However, chiral angle increased monotonically with temperature up to 42 °C (Fig. 5C), indicating that chirality is regulated by a factor other than growth rate that changes with temperature.

**Figure 5:**
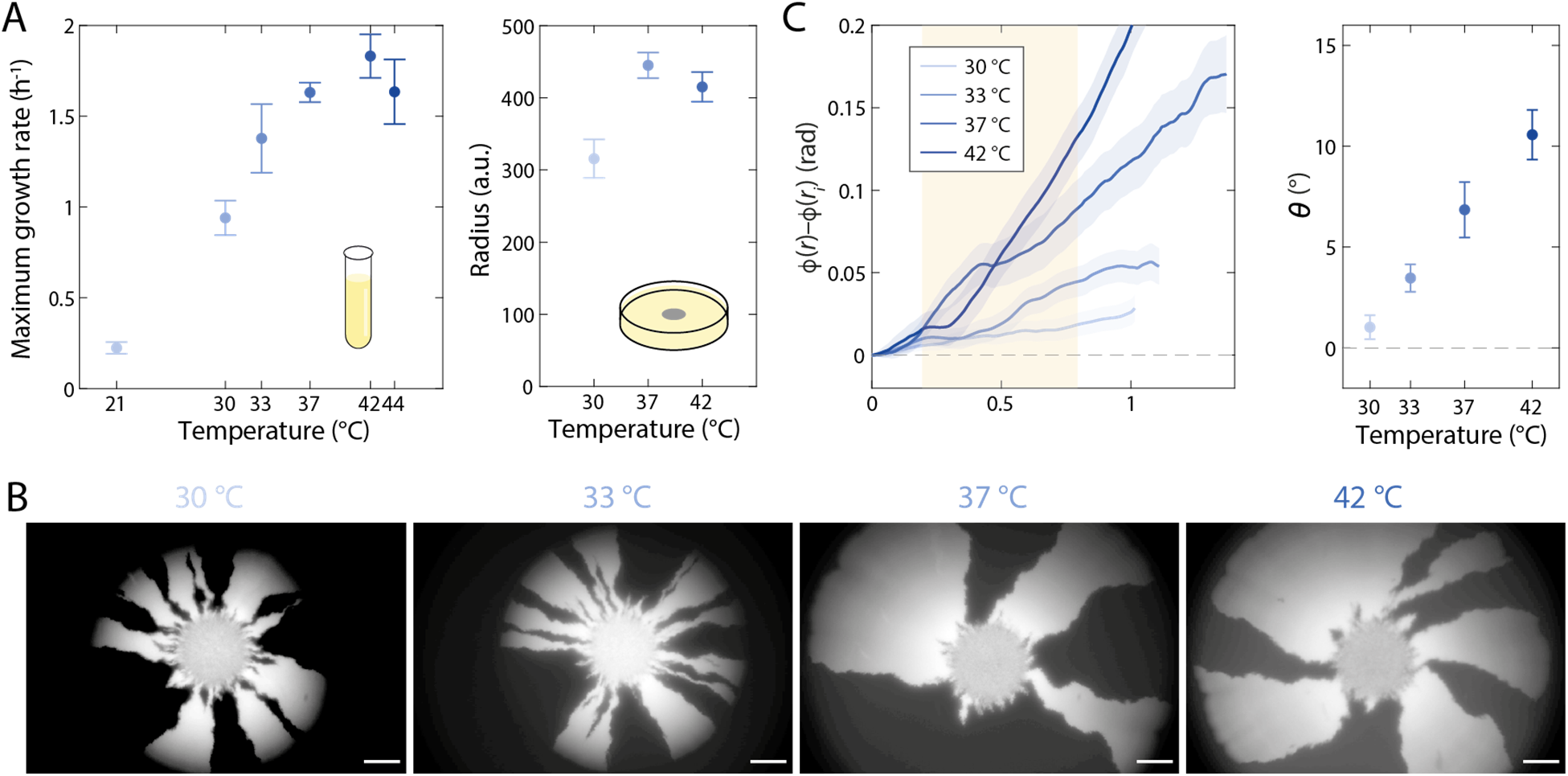
Chiral angle increases with increasing temperature. A) *E. coli* DH5*α* Growth depends on temperature. Left: maximum growth rate in liquid peaks at 42 °C. Right: Colony radius is higher at 37 °C than at 30 or 42 °C. B) Images of representative colonies show increasing chirality at higher temperatures. Scale bars: 1 mm. C) Left: mean rotation of sector boundaries at each temperature. Right: the chiral angle continues to increase with increasing temperature, even at 42 °C. Each data point is the average of *n*≥5 colonies. Error bars represent 1 standard deviation.

### Filamentation is linked to the extent of colony chirality

Single-cell twisting was approximately constant across temperatures (Fig. S5), again highlighting that another factor must be dictating chirality. Motivated by our observations that A22 treatment reduced cell length (Fig. 1D) and chiral angle (Fig. 2C), we measured log-phase DH5*α* cellular dimensions in liquid culture at various temperatures. Length and width both increased monotonically with temperature (26), with cells at 44 °C more than twice as long as those at 21 °C (Fig. S6). To verify whether cells were similarly elongated in colonies *in situ*, we imaged colonies directly and identified filamentous cells at the extreme edge of colonies with pronounced chirality (37 °C and 42 °C) (Fig. 6A), while less filamentation was apparent in colonies with little chirality (lower temperatures and anerobic growth) (Fig. 6 S7). Because of the difficulty in segmenting individual cells from *in situ* images of the colony, we sampled and imaged cells from the colony edge. We surmised that higher cell density in images was indicative of more sampling towards the center of the colony. Thus, to correct for variability in sampling location we focused our analysis on sets of images with similar cell density (Fig. S8). As in liquid, mean cell length increased with temperature and colonies grown at higher temperatures displayed more filamentous cells (Fig. 6B). Interestingly, cells grown in liquid had similar cellular dimensions aerobically and anaerobically, but cells sampled from the edge of colonies grown anaerobically were significantly shorter and wider than those from the edge of aerobic colonies (Fig. 6C, S9). These observations are consistent with the hypothesis that some degree of cell filamentation is necessary for colony chirality.

**Figure 6:**
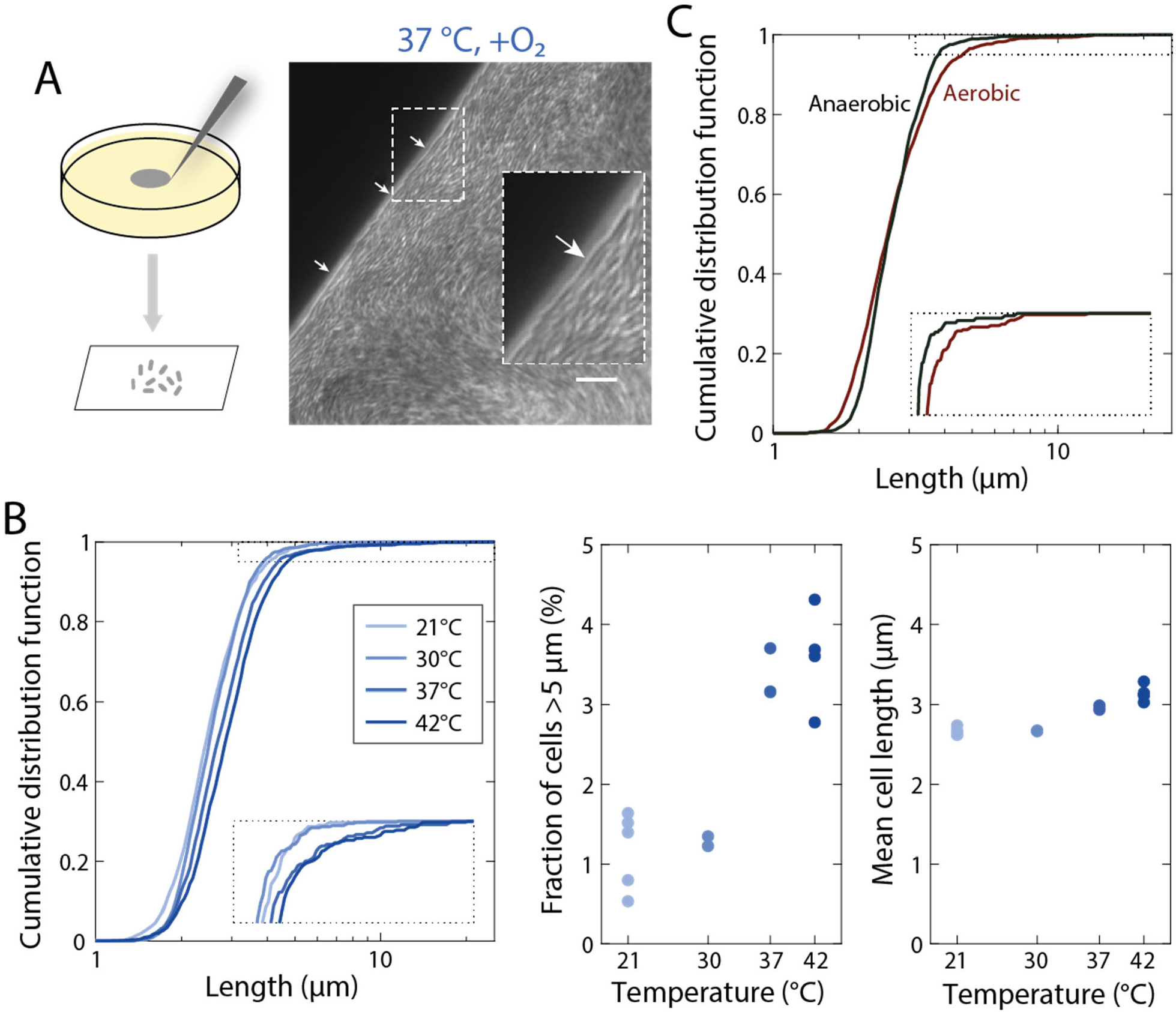
Conditions with enhanced colony chirality exhibit increased fractions of filamentous cells at the colony edge. A) Left: schematic of sampling from the colony edge. Right: imaging of the colony edge reveals filamentous cells at the border (arrows). Inset is a 200% zoom-in of the region surrounded by the dashed white line highlighting a filamentous cell. B) At higher temperatures, a larger fraction of the population exhibits filamentation. Left: the cumulative distribution function of cell length shifts to the right at increasing temperature. The inset is a zoom-in of the dotted region. Middle: the fraction of cells with length >5 µm in samples from the edge of colonies increases at increasing temperature. Right: mean cell length also increases slightly with increasing temperature. Each circle was computed from ≥16 fields of view from a sample from a distinct colony. C) During aerobic growth, a larger fraction of the population exhibits filamentation compared with anaerobic growth. The inset is a zoom-in of the dotted region.

### Inhibition of cell division enhances chirality

To determine whether filamentation and chirality are causally linked, we sought to increase the fraction of filamentous cells within a colony. We grew mixed cultures at 37 °C on various concentrations of cephalexin, a beta-lactam antibiotic that inhibits the division-specific transpeptidase PBP3 (27). At higher concentrations, colonies exhibited pronounced chirality similar to 42 °C, and chiral angle increased in a dose-dependent manner (Fig. 7A, S10). To test whether cephalexin would introduce chirality in situations where the chiral angle was close to zero, we grew mixed cultures anaerobically on LB+10 µg/mL cephalexin plates. Remarkably, cephalexin-treated colonies exhibited obvious chirality (Fig. 7C). Thus, inducing filamentation is sufficient to introduce or accentuate colony chirality.

**Figure 7:**
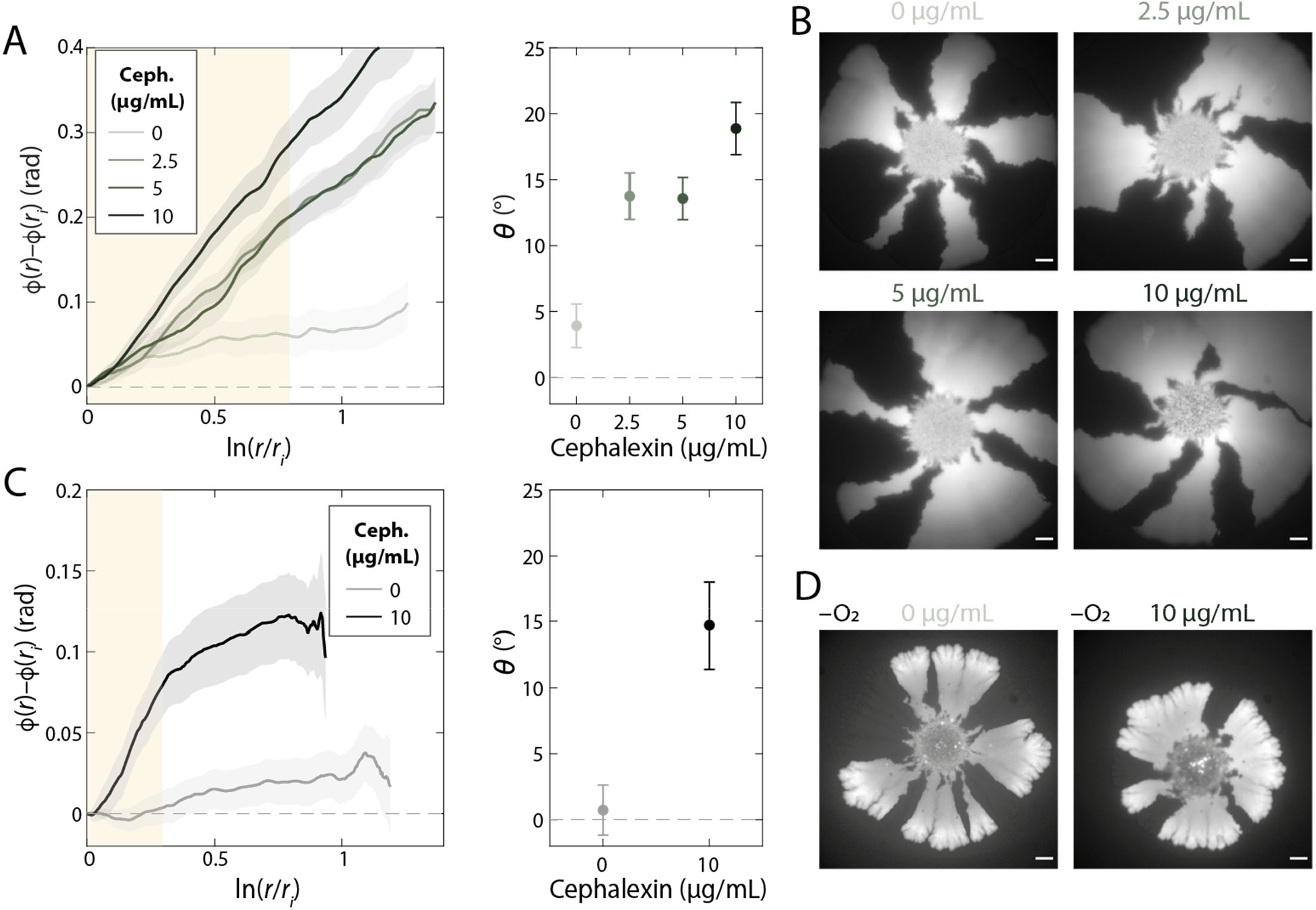
Division inhibition using cephalexin results in enhanced chirality during aerobic growth and the introduction of chirality during anaerobic growth. A) Left: mean rotation of sector boundaries of aerobically grown DH5*α* wild-type colonies with various concentrations of cephalexin. Right: the chiral angle increases with increasing cephalexin concentration. Each data point is the average of *n*≥5 colonies. Error bars represent 1 standard deviation. B) Representative colonies grown aerobically with various concentrations of cephalexin. Scale bars: 1 mm. C) Left: mean rotation of sector boundaries of anaerobically grown DH5*α* wild-type colonies without and with cephalexin. Right: in the presence, but not in the absence, of cephalexin colonies exhibit chirality. Each data point is the average of *n*≥5 colonies. Error bars represent 1 standard deviation. D) Representative colonies grown anaerobically without and with 10 µg/mL cephalexin. Scale bars: 1 mm.

## Discussion

In this study, we systematically measured the degree and handedness of twisting at the single-cell level across many perturbations, and determined that while changes that perturb single-cell twisting alter colony chirality, changes that perturb colony chirality do not necessarily alter single-cell twisting. It is clear that many factors interact to determine colony structure, including forces between cells and accessibility of nutrients (28), cellular geometry (29), and interactions of cells with each other and with the surface (20). Previous findings connecting surface attachment with colony chirality (20) are not inconsistent with our study, since the dependence of agar properties and surface attachment on temperature, cell size, and oxygen are unknown. Our colony chirality observations are phenomena that are either intrinsically three-dimensional or originate from cells being subject to a solid surface on one side and an air surface on the other, as confining the cells between two agar pads may eliminate chirality by removing the air-agar asymmetry (Fig. 4A). Regardless, our data highlight the role of cell filamentation in establishing colony chirality, with division-inhibiting cephalexin treatment leading to the introduction of chirality in colonies grown anaerobically (Fig. 7). Moreover, cell length is the variable that unifies colony chirality across all of our growth conditions. These results reveal a complex connection between single-cell dimensions and population level spatial patterns (30, 31).

Our original motivation for this study was to establish whether colony chirality can be traced back to single-cell behaviors. While we have shown that cell filamentation is coupled to the generation of colony chirality, it remains unclear whether single-cell twisting is also a factor. A22 treatment does reduce chirality (Fig. 2) as does genetic manipulation of the cell-wall synthesis machinery (Fig. 3). Moreover, single-cell twisting and colony chirality each have a defined handedness in the absence of A22. However, it remains unclear whether and how handedness at the microscopic scale determines handedness at the macroscopic scales.

The connection between filamentation, and more generally cell shape, and colony patterning was unexpected, in part because the regulation of division across environmental conditions such as A22 treatment, anaerobic growth, and temperature are poorly understood. In the case of A22, it was critical to measure cell shape at the edge of the colony, which revealed that the cell width phenotype characteristic of A22 treatment on short time scales was reversed after several days (Fig. 2D), likely due to transcriptional adaptation (21). It is intriguing that regulation of cell division takes place on such long time scales. The tunability of chirality using cephalexin provides an interesting control knob for the design of macroscale patterns in bacterial communities.

What are the origins of colony chirality? Ultimately, chirality must materialize from some molecular symmetry breaking, and while the cell wall is an enticing candidate, the extracellular environment cannot be ruled out. This knowledge gap motivates the quantification of single-cell twisting and colony chirality across more strains and species, to determine whether the two are generally coupled. An ideal demonstration would be the reversal of handedness of a single species, as we attempted in this work, although it may be impossible to construct a right-handed *E. coli* cell that is stable during filamentation without reprogramming many cell-wall synthesis components.

A wealth of exotic patterns and mechanical properties can emerge from chiral constituents exerting forces on each other and their environments (32-34), including periodic crystals of rotating particles (35-37) that can synchronize into exotic phases (38). Mixtures of oppositely rotating particles tend to phase separate (39, 40), leading to complex structures and flows at chiral-phase interfaces (41, 42). Analyzing flow patterns and excitations of chiral active fluids has led to the design of topological states of matter (43, 44) and predictions of novel hydrodynamic responses (45, 46) that have been experimentally measured (47). Biological settings such as colony chirality may provide even more complex ways to connect microscopic forces of chiral active components to macroscopic handedness. For example, the observation that filamentation in only a few percent of cells is sufficient to induce chirality in anaerobically grown colonies motivates the study of mixtures of active particle with a heterogeneous distribution of geometries.

From an ecological and evolutionary perspective, what is the relevance of colony chirality? Recent work has suggested that chirality is an important population-level trait that mediates competition, invasion, and ultimately spatial structure within a bacterial community (48). If so, then our work suggests that colony-scale patterning has likely applied selective pressures on division at the cellular-scale. To design the spatial structure of bacterial communities, it is advantageous to control the emergent pattern of cell growth rather than have to establish patterns from the start. To achieve this control will ultimately require a mechanistic model for colony chirality, the development of which will be facilitated by the discovery that cellular properties such as cell length are critical parameters.

## Materials and Methods

### Bacterial strains and plasmids

Strains used in this study are described in Table S1. In brief, we used three pairs of strains. In each pair, the two strains have a common genetic background and carry plasmids expressing CFP or venus YFP. The pair used for the majority of experiments is identical to the one used in previous studies reporting colony chirality (17, 19, 20). In each pair, there were no obvious fitness differences between the two strains (both strains were able to form sectors). Fluorescence levels were sufficient for imaging even for basal expression in the absence of IPTG, hence we did not include IPTG in any of the media.

For this study, we constructed *E. coli* DH5*α* Δ*mrdA* strains with constitutive expression of *V. cholerae mrdA* by replacing the genomic *E. coli mrdA* sequence with a 4169-bp PCR fragment encoding the *V. cholerae mrdA* (Vc-*mrdA*) sequence and kanamycin resistance cassette, followed by selection for stability. Construction of the DH5*α* Δ*mrdA* strain was as follows: the Vc-*mrdA* DNA fragment was amplified from the pww308-VcmrdA construct (15) with primers that have 20 nucleotides of homology to the plasmid DNA sequence and 50 nucleotides of homology to the genomic Ec-*mrdA* target sequence, specifically delmrdAFor (5’-ACGCAGCGGATGAAACTACAGAACTCTTTTCGCGACTATACGGCAGAGCCACGTTGTGTCTCAAA ATC-3’) and delmrdARev (5’-TATCCGTCATGATTAATGGTCCTCCGCTGCGGCAACCGCTGGATTTTCCGCATCCTAGGTCATGGCTGTATTAC). The DH5*α* strain lacks the ability for homologous recombination. Therefore, *E*.*coli* strain DH5*α* (Invitrogen) was first transformed with the pSIM5 plasmid (pSC101 repA^ts^) encoding Red recombination proteins. Selection of transformed cells was performed by growing in Lennox LB in the presence of 17.5 µg/mL chloramphenicol at 30°C to allow pSIM5 replication, producing strain eCR106 that was able to perform homologous recombination. eCR106 cells were transformed with the gel-purified Vc-*mrdA* DNA fragment via electroporation as follows: eCR106 cells were grown at 30 °C to OD=0.4-0.5, transferred to 42 °C for 15 min for induction of Red proteins, and then transferred to ice for 10 min. Cells were centrifuged at 3000*g* and the pellet was washed with ice-cold water twice before resuspending in 10% glycerol solution for electroporation using a GenePulser XCell (BioRad) electroporator with the pre-set bacterial protocol for *E. coli* using a 1-mm cuvette. After transformation, cells were incubated at 30 °C for 2 h before selection on Lennox LB agar plates with 25 μg/mL kanamycin. Transformed cells were identified by colony PCR with primers delBFor (5’–ACATCATCGCCTTAGACGTTC-3’) and delBRev (5’-AGATGGACTTTATCCCAGAATG-3’) for the upstream/downstream *E. coli mrdA* sequence and primers delCFor (5’-AGCGGATGAAACTACAGAACTC-3’) and delCRev (5’-CGGCAACCGCTGGATTTTC).

Colonies were picked and grown in Lennox LB at 37 °C to remove the pSIM5 plasmid, creating strain eCR111. pSIM5 loss was confirmed by the absence of growth under chloramphenicol selection. Genomic eCR111 DNA was extracted using a GeneJET Genomic DNA Purification Kit (ThermoFisher) and used as DNA template for amplification with primers delBFor and delCRev. The PCR fragment was sequenced using a MinION (Oxford Nanopore Technologies). Sequencing results showed that the Tet promoter region of the Vc-*mrdA* fragment used for the recombination was lost in this strain.

CFP/YFP plasmids were constructed by cloning the fluorescent protein coding sequence into pSTV28 (Takara Bio Europe SAS) with EcoRI/SalI restriction sites. eCR111 cells were transformed with CFP/YFP constructs via electroporation as above, generating strains eCR112_1 with CFP and eCR112_2 with YFP. Selection of transformants was achieved using 35 µg/mL chloramphenicol.

### Growth of mixed colonies

A step-by-step protocol is available online on protocols.io (https://www.protocols.io/view/growth-of-mixed-e-coli-colonies-bqgjmtun). In short, agar plates (6 cm in diameter) were prepared with 10 mL of Lennox LB (RPI, Cat. #L24066; Melford, Cat. #L24060-100.0 for colonies sandwiched between agar slabs and corresponding control experiments) with 1.5% agar (BD, Cat. #214530) and left on the bench overnight. Plates were used the next day or stored at 4 °C. As necessary, A22 or cephalexin were added from frozen stocks to the liquid after autoclaving. To initiate colony growth, the appropriate pair of fluorescent *E. coli* strains were grown overnight at 37 °C in liquid Lennox LB. Both cultures were diluted 10-fold in fresh LB and mixed. One microliter droplets were pipetted onto prewarmed plates and grown at 37 °C, typically for 7 to 8 days. Anaerobic growth experiments were carried out in a custom anaerobic chamber (Coy Lab Products). Colony growth between agar surfaces was achieved by cutting an agar pad out of a fresh plate and placing it upside down onto freshly inoculated cultures.

### Sampling from mixed colonies

Cells were sampled from the border of colonies using a 20-µL LTS micropipette tip (Rainin). After touching the edge of the colony, cells were resuspended in 20 µL PBS and 1 µL was spotted onto PBS 1% agarose pads for imaging.

### Imaging of mixed colonies

Fluorescence images of colonies were acquired after 7-8 days of growth using a 1X objective on a Nikon Eclipse Ti-E inverted fluorescence microscope equipped with a DU897 electron multiplying charged couple device (EMCCD) camera (Andor) using µManager v. 1.4 (49), or a Nikon TE-2000 or Zeiss Axio Zoom.V16 microscope. Colonies sandwiched between two agar surfaces and the colonies in the corresponding control experiments were imaged after 7 days at 4 °C. Colonies that did not show sufficient fluorescence at this point, were imaged again after 26 more days at 4 °C.

Edges of colonies were imaged with a 20X objective on a Nikon Eclipse Ti-E inverted fluorescence microscope equipped with a DU897 camera (Andor) using µManager v. 1.4.

### Identifying colony radius and sector boundaries in mixed colonies

Images were imported into Matlab. The difference between YFP and CFP images was used to identify boundaries of intensity using the ‘edge’ function with the ‘Log’ method. The center of the colony was defined by fitting a circle to 20 points manually selected from the images either from the border of the colony if the colonies were small enough to fit in the image, or from the border of the mixed-fluroescence sector at the center of the colony. The coordinates of the boundaries were transformed to polar coordinates based on the center of the colony. Points were mapped to traces by connecting nearest neighbors. Points too close to the center of the colony were discarded. Boundary traces were cleaned up manually: some were separated at a manually selected location when the traces clearly represented two sides of a sector, and some were removed because they clearly captured a feature that was not a sector boundary. Traces in polar coordinates were then smoothed. Some parameters (closeness to the center or the threshold for edge detected) were modified depending on the quality of the image. All traces from all colonies of a given experiment were then pooled. The mean was obtained by bootstrapping: random traces (selected with replacement) were averaged to calculate the slope, and then the slope was integrated to obtain a mean trace. This process was repeated 200 times to compute the final mean trace and the standard deviation over the 200 iterations.

All code is available in a GitHub repository https://github.com/aarandad/ColonyChirality-KCH.

### Growth measurements in liquid culture

Optical density (OD) measurements were performed with an Epoch 2 plate reader (BioTek) at 37 °C with continuous shaking and OD_600_ measured at 7.5-min intervals. Maximal growth rate was calculated as the maximal slope of ln(OD) with respect to time (calculated from a linear regression of a sliding window of 11 timepoints) using custom Matlab R2018b (Mathworks) code.

### Single-cell imaging

Cultures were grown overnight at 37 °C in LB, and diluted 1:200 into fresh medium (with antibiotics when appropriate). For phase-contrast imaging, cells were spotted onto a 1% (w/v) agarose pad with LB. Cells treated with A22 were exposed to the drug for 2-3 h before imaging.

Phase-contrast and epifluorescence images were acquired with a Nikon Ti-E inverted microscope (Nikon Instruments) using a 100X (NA 1.40) oil immersion objective and a Neo 5.5 sCMOS camera or a DU885 EMCCD (Andor Technology). The microscope was outfitted with an active-control environmental chamber for temperature regulation (HaisonTech, Taipei, Taiwan). Images were acquired using µManager v. 1.4 (49). Cell contours were automatically segmented using *Morphometrics* (50) and a local coordinate system was defined based on the meshing algorithm from *MicrobeTracker* (51). Some images of cells sampled from the edge of colonies (Fig. 6) had clusters of cells that could not be segmented by *Morphometrics*. These images were processed with the neural network-based machine learning framework *DeepCell* (52) prior to segmentation with *Morphometrics*.

### Twist ‘n’ TIRF measurements

Fluorescent WGA was added to liquid cultures to a final concentration of 25 µg/mL. After h of growth, cells were pelleted at 8,000*g* for 1 min and washed with PBS once before being spotted onto a 1% (w/v) EZ-RDM 0.2% glucose or LB agarose pads with 10 µg/mL cephalexin. For anaerobic Twist ‘n’ TIRF experiments, cultures were grown, washed, and spotted onto agarose pads inside an anaerobic chamber. Mounted pads were fully sealed with VALAP before removal from the anaerobic chamber and imaging was conducted immediately.

### TIRF microscopy

Twisting measurements were performed on a Ti-E inverted microscope (Nikon, NY, USA) with a 100X objective (NA 1.40). A Sapphire OPSL 488-nm laser (Coherent, CA, USA) was coupled into a TIRF illuminator (Nikon) attached to the microscope stand. Images were acquired with DU885 EMCCD (Andor, CT, USA) camera and synchronization between components was achieved using µManager (49).

### Cell twisting analysis

In Twist ‘n’ TIRF experiments, cell contours were computationally extracted from phase using Morphometrics (50). The integrated WGA fluorescence enclosed within the cell contour was quantified. These values were normalized to the pre-bleached level and plotted as a function of the change in length Δ*l* relative to the pre-bleached length. Curves were then fitted to extract the slope *λ* representing the rate of fluorescence recovery due to twisting. The curves were fit over the first 3 µm of elongation to avoid noise from photobleaching after large amounts of exposure.

## Acknowledgements

We thank members of the Huang and Hallastchek lab for helpful discussions. This work was funded by NIH CAREER Award MCB-1149328 (to K.C.H.), the Allen Discovery Center at Stanford University on Systems Modeling of Infection (to K.C.H.). A.A.-D. is a Howard Hughes Medical Institute International Student Research fellow, a Stanford Bio-X Bowes fellow, and a Siebel Scholar. K.C.H. is a Chan Zuckerberg Investigator.

## References

1. P. Satir, Chirality of the cytoskeleton in the origins of cellular asymmetry. Philos Trans R Soc Lond B Biol Sci 371 (2016).

2. H. Shi, D. A. Quint, G. M. Grason, A. Gopinathan, K. C. Huang, Chiral twisting in a bacterial cytoskeletal polymer affects filament size and orientation. Nat Commun 11, 1408 (2020).

3. S. Hozumi et al., An unconventional myosin in Drosophila reverses the default handedness in visceral organs. Nature 440, 798–802 (2006).

4. M. Inaki, J. Liu, K. Matsuno, Cell chirality: its origin and roles in left-right asymmetric development. Philos Trans R Soc Lond B Biol Sci 371 (2016).

5. S. R. Naganathan, S. Furthauer, M. Nishikawa, F. Julicher, S. W. Grill, Active torque generation by the actomyosin cell cortex drives left-right symmetry breaking. Elife 3, e04165 (2014).

6. T. Shoji et al., Plant-specific microtubule-associated protein SPIRAL2 is required for anisotropic growth in Arabidopsis. Plant Physiol 136, 3933–3944 (2004).

7. T. S. Ursell et al., Rod-like bacterial shape is maintained by feedback between cell curvature and cytoskeletal localization. Proceedings of the National Academy of Sciences of the United States of America 111, E1025–1034 (2014).

8. S. Wang, L. Furchtgott, K. C. Huang, J. W. Shaevitz, Helical insertion of peptidoglycan produces chiral ordering of the bacterial cell wall. Proceedings of the National Academy of Sciences of the United States of America 109, E595–604 (2012).

9. N. H. Mendelson, Helical growth of Bacillus subtilis: a new model of cell growth. Proceedings of the National Academy of Sciences of the United States of America 73, 1740–1744 (1976).

10. J. V. Holtje, Growth of the stress-bearing and shape-maintaining murein sacculus of Escherichia coli. Microbiol Mol Biol Rev 62, 181–203 (1998).

11. K. D. Young, The selective value of bacterial shape. Microbiol Mol Biol Rev 70, 660–703 (2006).

12. L. Furchtgott, N. S. Wingreen, K. C. Huang, Mechanisms for maintaining cell shape in rod-shaped Gram-negative bacteria. Mol Microbiol 81, 340–353 (2011).

13. Z. Gitai, N. Dye, L. Shapiro, An actin-like gene can determine cell polarity in bacteria. Proceedings of the National Academy of Sciences of the United States of America 101, 8643–8648 (2004).

14. Z. Gitai, N. A. Dye, A. Reisenauer, M. Wachi, L. Shapiro, MreB actin-mediated segregation of a specific region of a bacterial chromosome. Cell 120, 329–341 (2005).

15. C. Tropini et al., Principles of bacterial cell-size determination revealed by cell-wall synthesis perturbations. Cell Rep 9, 1520–1527 (2014).

16. S. van Teeffelen et al., The bacterial actin MreB rotates, and rotation depends on cell-wall assembly. Proceedings of the National Academy of Sciences of the United States of America 108, 15822–15827 (2011).

17. O. Hallatschek, P. Hersen, S. Ramanathan, D. R. Nelson, Genetic drift at expanding frontiers promotes gene segregation. Proceedings of the National Academy of Sciences of the United States of America 104, 19926–19930 (2007).

18. O. Hallatschek, D. R. Nelson, Gene surfing in expanding populations. Theor Popul Biol 73, 158–170 (2008).

19. K. S. Korolev, J. B. Xavier, D. R. Nelson, K. R. Foster, A quantitative test of population genetics using spatiogenetic patterns in bacterial colonies. Am Nat 178, 538–552 (2011).

20. L. Jauffred, R. Munk Vejborg, K. S. Korolev, S. Brown, L. B. Oddershede, Chirality in microbial biofilms is mediated by close interactions between the cell surface and the substratum. ISME J 11, 1688–1701 (2017).

21. M. Silvis et al., Morphological and transcriptional responses to essential-gene depletion in Escherichia coli. (2021).

22. B. G. Spratt, Distinct penicillin binding proteins involved in the division, elongation, and shape of Escherichia coli K12. Proceedings of the National Academy of Sciences 72, 2999–3003 (1975).

23. S. J. Remington, Fluorescent proteins: maturation, photochemistry and photophysics. Curr Opin Struct Biol 16, 714–721 (2006).

24. R. Y. Tsien, The green fluorescent protein. Annu Rev Biochem 67, 509–544 (1998).

25. M. A. Barber, The Rate of Multiplication of Bacillus Coli at Different Temperatures. The Journal of Infectious Diseases 5, 379–400 (1908).

26. M. Schaechter, O. Maaløe, N. Kjeldgaard, Dependency on medium and temperature of cell size and chemical composition during balanced grown of Salmonella typhimurium. J. Gen. Microbiol. 19, 592–606 (1958).

27. P. J. Hedge, B. G. Spratt, Amino acid substitutions that reduce the affinity of penicillin-binding protein 3 of Escherichia coli for cephalexin. Eur J Biochem 151, 111–121 (1985).

28. M. R. Warren et al., Spatiotemporal establishment of dense bacterial colonies growing on hard agar. Elife 8 (2019).

29. W. P. Smith et al., Cell morphology drives spatial patterning in microbial communities. Proceedings of the National Academy of Sciences of the United States of America 114, E280–E286 (2017).

30. J. van Gestel, F. J. Weissing, O. P. Kuipers, A. T. Kovacs, Density of founder cells affects spatial pattern formation and cooperation in Bacillus subtilis biofilms. ISME J 8, 2069–2079 (2014).

31. E. Ben-Jacob, I. Cohen, H. Levine, Cooperative self-organization of microorganisms. Advances in Physics 49, 395–554 (2000).

32. J. C. Tsai, F. Ye, J. Rodriguez, J. P. Gollub, T. C. Lubensky, A Chiral Granular Gas. Phys. Rev. Lett. 94, 214301 (2005).

33. S. Fürthauer, M. Strempel, S. W. Grill, F. Jülicher, Active chiral fluids. Eur. Phys. J. E 35, 89 (2012).

34. K. Drescher et al., Dancing <i>Volvox</i> : Hydrodynamic Bound States of Swimming Algae. Phys. Rev. Lett. 102, 168101 (2009).

35. A. P. Petroff, X.-L. Wu, A. Libchaber, Fast-Moving Bacteria Self-Organize into Active Two-Dimensional Crystals of Rotating Cells. Phys. Rev. Lett. 114, 158102 (2015).

36. J. Yan, S. C. Bae, S. Granick, Rotating crystals of magnetic Janus colloids. Soft Matter 11, 147–153 (2015).

37. A. Snezhko, Complex collective dynamics of active torque-driven colloids at interfaces. Current Opinion in Colloid & Interface Science 21, 65–75 (2016).

38. B. C. van Zuiden, J. Paulose, W. T. M. Irvine, D. Bartolo, V. Vitelli, Spatiotemporal order and emergent edge currents in active spinner materials. Proceedings of the National Academy of Sciences 113, 12919–12924 (2016).

39. N. H. P. Nguyen, D. Klotsa, M. Engel, S. C. Glotzer, Emergent Collective Phenomena in a Mixture of Hard Shapes through Active Rotation. Phys. Rev. Lett. 112, 075701 (2014).

40. K. Yeo, E. Lushi, P. M. Vlahovska, Collective Dynamics in a Binary Mixture of Hydrodynamically Coupled Microrotors. Phys. Rev. Lett. 114, 188301 (2015).

41. C. del Junco, L. Tociu, S. Vaikuntanathan, Energy dissipation and fluctuations in a driven liquid. Proceedings of the National Academy of Sciences 115, 3569–3574 (2018).

42. C. Scholz, M. Engel, T. Pöschel, Rotating robots move collectively and self-organize. Nature Communications 9, 931 (2018).

43. A. Souslov, K. Dasbiswas, M. Fruchart, S. Vaikuntanathan, V. Vitelli, Topological Waves in Fluids with Odd Viscosity. Phys. Rev. Lett. 122, 128001 (2019).

44. G. Baardink, G. Cassella, L. Neville, P. A. Milewski, A. Souslov, Complete absorption of topologically protected waves. arXiv:2010.07342 [cond-mat, physics:physics] (2020).

45. D. Banerjee, A. Souslov, A. G. Abanov, V. Vitelli, Odd viscosity in chiral active fluids. Nature Communications 8, 1573 (2017).

46. A. Souslov, A. Gromov, V. Vitelli, Anisotropic odd viscosity via a time-modulated drive. Phys. Rev. E 101, 052606 (2020).

47. V. Soni et al., The odd free surface flows of a colloidal chiral fluid. Nat. Phys. 15, 1188–1194 (2019).

48. B.G.A K. S. Korolev, Chirality provides a direct fitness advantage and facilitates intermixing in cellular aggregates. PLoS Comput Biol 14, e1006645 (2018).

49. N. Stuurman, N. Amdodaj, R. Vale, μManager: open source software for light microscope imaging. Microscopy Today 15, 42–43 (2007).

50. T. Ursell et al., Rapid, precise quantification of bacterial cellular dimensions across a genomic-scale knockout library. BMC Biol 15, 17 (2017).

51. O. Sliusarenko, J. Heinritz, T. Emonet, C. Jacobs-Wagner, High-throughput, subpixel precision analysis of bacterial morphogenesis and intracellular spatio-temporal dynamics. Molecular microbiology 80, 612–627 (2011).

52. D. A. Van Valen et al., Deep Learning Automates the Quantitative Analysis of Individual Cells in Live-Cell Imaging Experiments. PLoS Comput Biol 12, e1005177 (2016).

