## Supplemental Tables and Figures for "Bacterial filamentation drives colony chirality"

**Table S1: Bacterial strains used in this study.**

| **Strain name** | **Relevant characteristics** | **Reference/origin** |
|---|---|---|
| DH5α-H-CFP | Plasmid pTrc99A with gene coding for CFP | (17) |
| DH5α-H-YFP | Plasmid pTrc99A with gene coding for YFP | (17) |
| DH5α-E-CFP | Plasmid pSTV28 with gene coding for CFP | This study |
| DH5α-E-YFP | Plasmid pSTV28 with gene coding for YFP | This study |
| DH5α-E-CFP Vc-*mrdA* | Plasmid as above, ΔEc-*mrdA* | This study |
| DH5α-E-YFP Vc-*mrdA* | Plasmid as above, ΔEc-*mrdA* | This study |

### Supplemental Figures

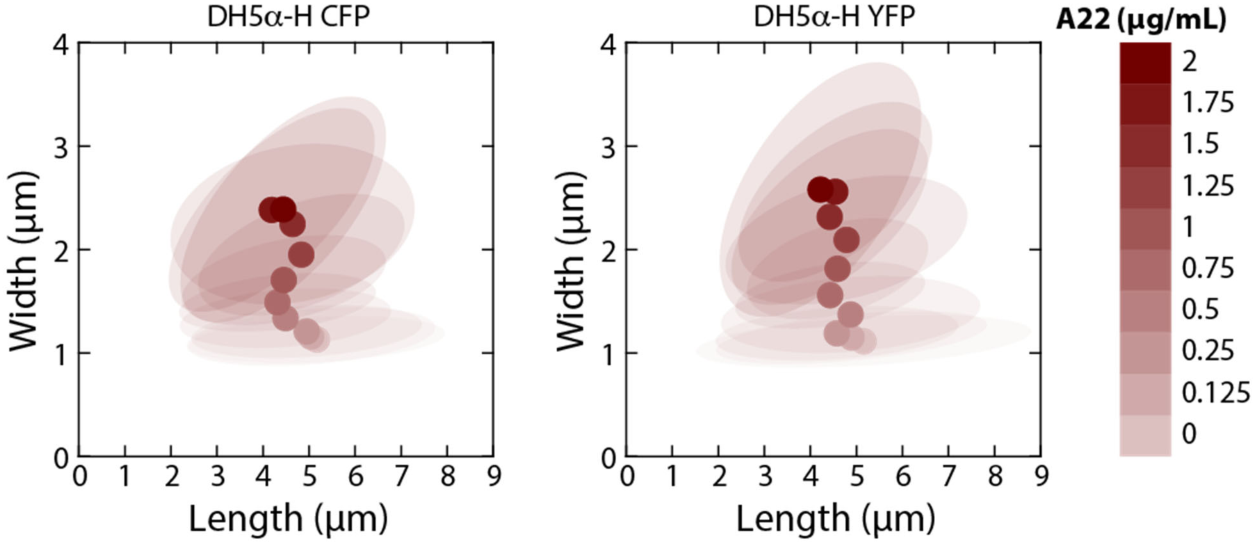


**Figure S1: Fluorescent protein-expressing strains have similar responses to A22 treatment.**

During liquid growth, the cell width of strains 282 and 284 expressing CFP and YFP, respectively, measured during log-phase growth in liquid increased as A22 concentration increased. Cell length decreased with increasing A22 concentration up to 1 μg/mL. Circles represent mean dimensions and ellipses represent the covariance matrix of length and width. *n*>50 cells were quantified for each condition.

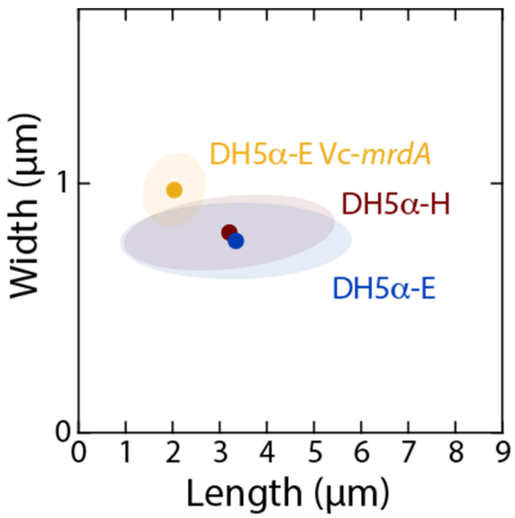


**Figure S2: *E. coli* DH5α-E Δ*mrdA* cells with heterologous expression of the *mrdA* gene from *Vibrio cholerae* (Vc-*mrdA*) remain wider than wild-type DH5α-H or DH5α-E cells after 5 days of colony growth*.***

Circles represent mean values and ellipses represent the covariance of width and length. *n*>50 cells were quantified for each strain.

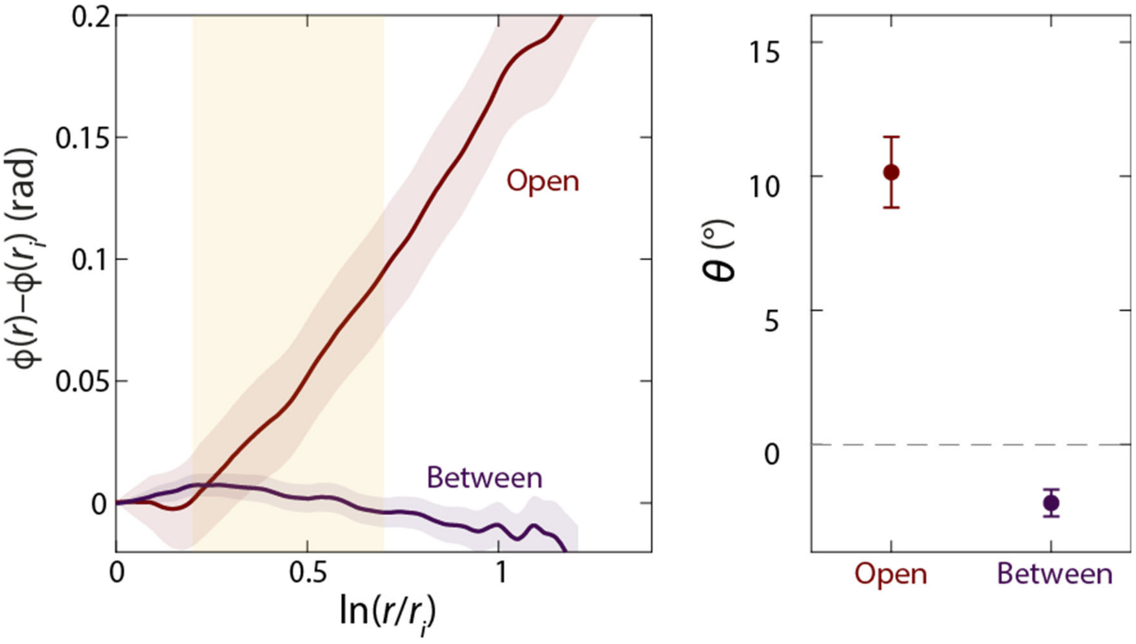


**Figure S3: Colony chirality is reduced when grown sandwiched between two agar surfaces.**

Shown is a biological replicate of the experiment in Fig. 4B. Left: mean rotation of sector boundaries in open or sandwiched configurations. Right: the chiral angle was zero or slightly negative for sandwiched colonies. Each data point is the average of $n \geq 5$ colonies. Error bars represent 1 standard deviation.

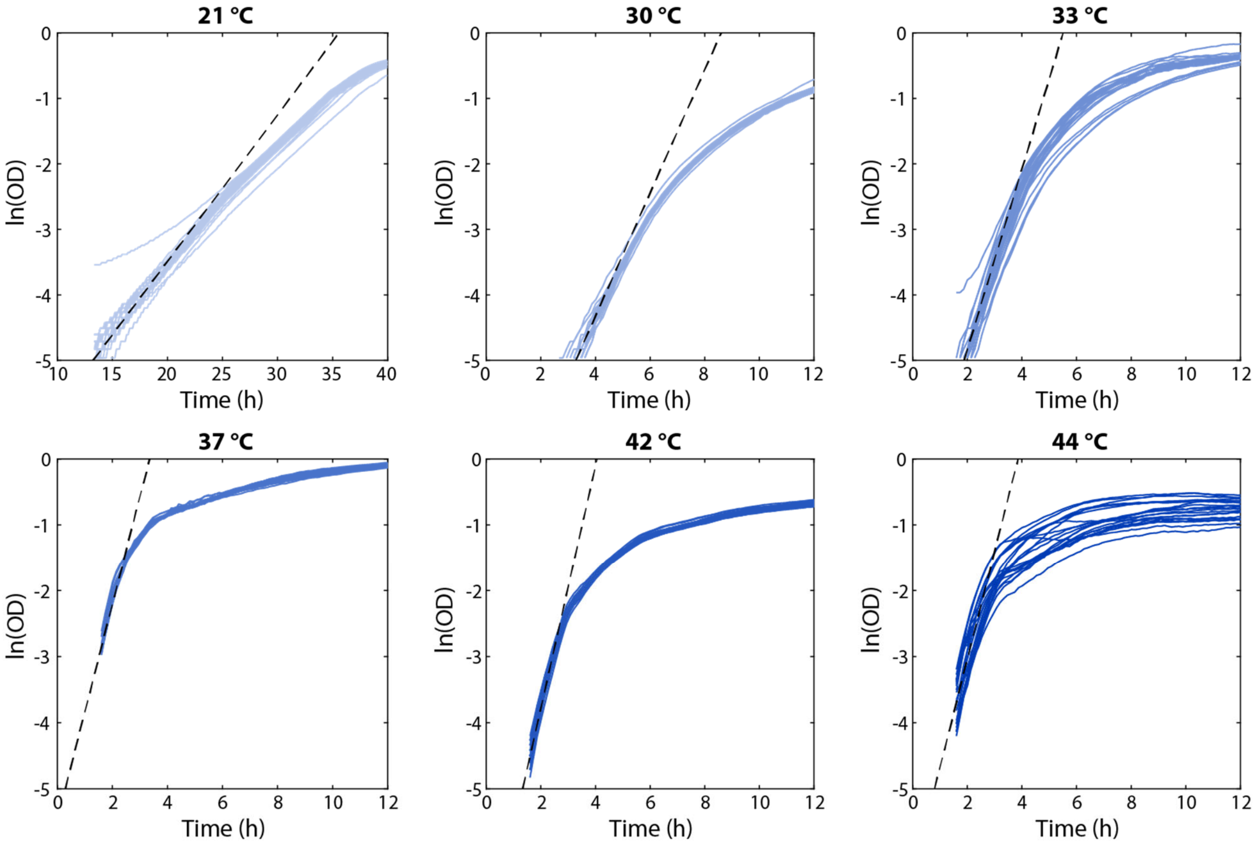


**Figure S4: Growth curves of *E. coli* DH5$\alpha$ grown in liquid at various temperatures.**

The slope of the dashed lines defined the maximum growth rate in Fig. 5A. Each plot shows 20 replicate growth curves.

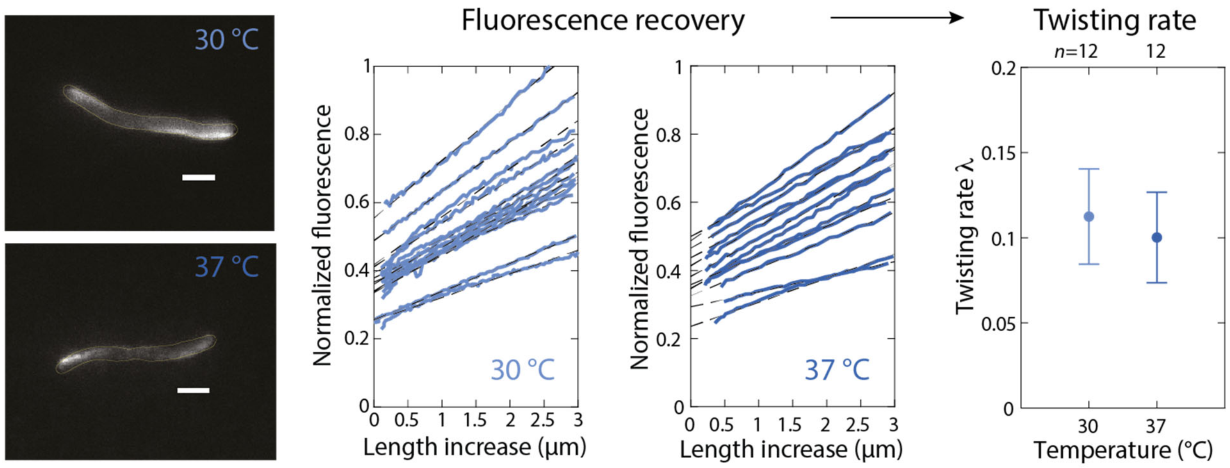


**Figure S5: The twisting rate is similar across temperatures.**

Fluorescence recovery (middle) after photobleaching of the bottom surface during Twist-n-TIRF experiments (left) was used to estimate the twisting rate (right) (Methods). Scale bars: 2 μm. All cells whose handedness could be reliably classified were left-handed (*n*=67, 30 °C; *n*=43, 37 °C).

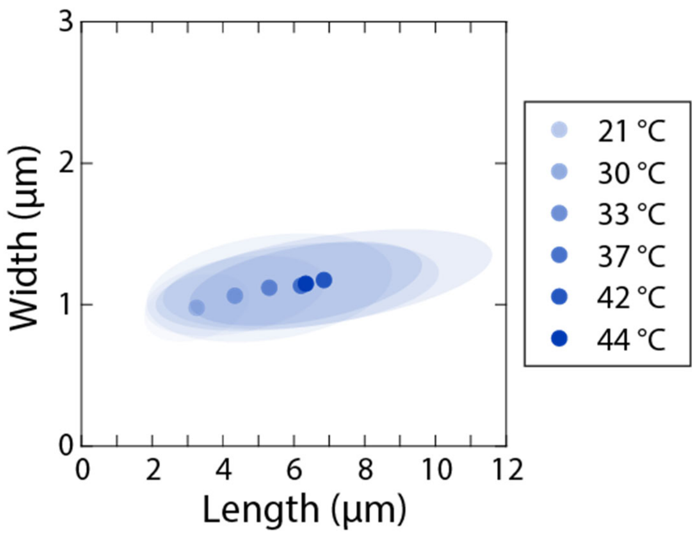


**Figure S6: Cell length generally increases with increasing temperature.**

During liquid growth, cell length measured during log-phase growth in liquid increased as temperature increased up to 42 °C. Circles represent mean dimensions and ellipses represent the covariance matrix of length and width. *n*>50 cells were quantified for each condition.

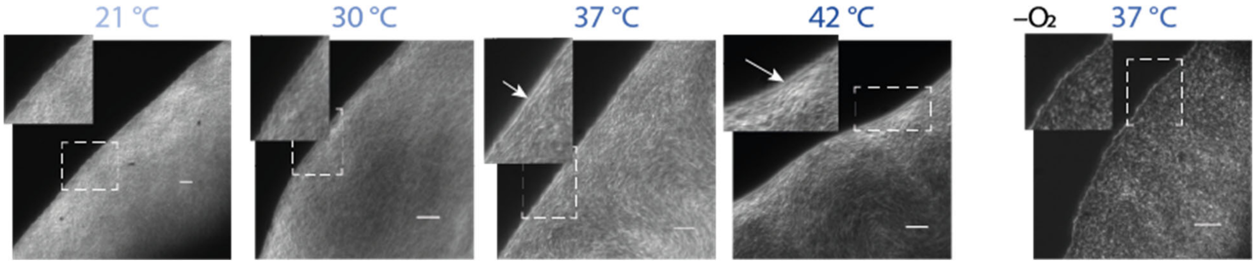


**Figure S7: Colonies grown at high temperatures have more filamentous cells at the border than colonies grown at low temperatures or in anaerobic conditions ($-O_2$).** Insets: zoom-ins of regions highlighted by white dashed line by 160%. Arrows denote filamentous cells. Scale bars: 10 μm.

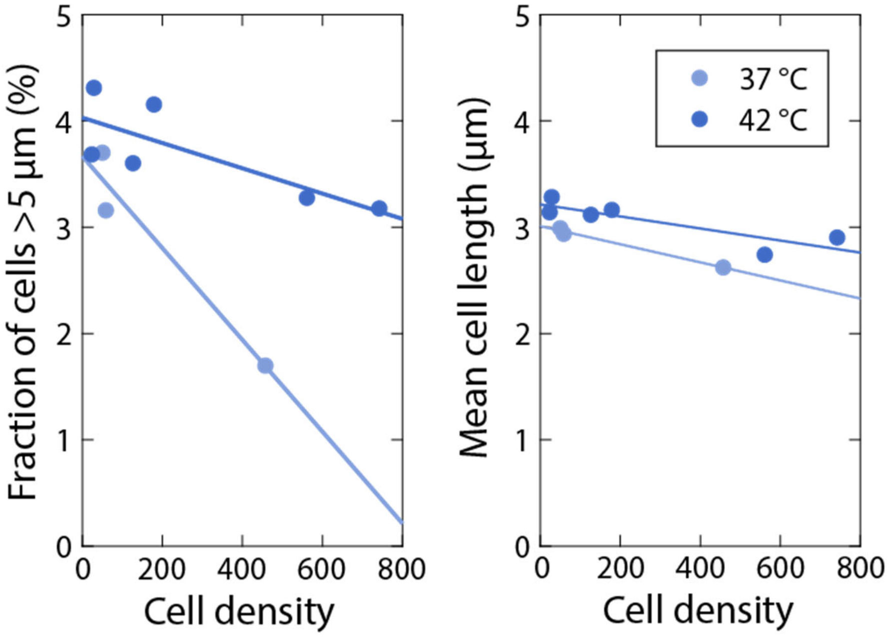


**Figure S8: Cell length varies with cell density.**

Cells were sampled from the edge of colonies grown at 37 or 42 °C. For samples with a higher cell density (likely due to sampling farther from the colony edge), the fraction of cells longer than 5 μm (left) and the mean cell length (right) was lower. Each circle was computed from ≥16 fields of view from a sample from a distinct colony.

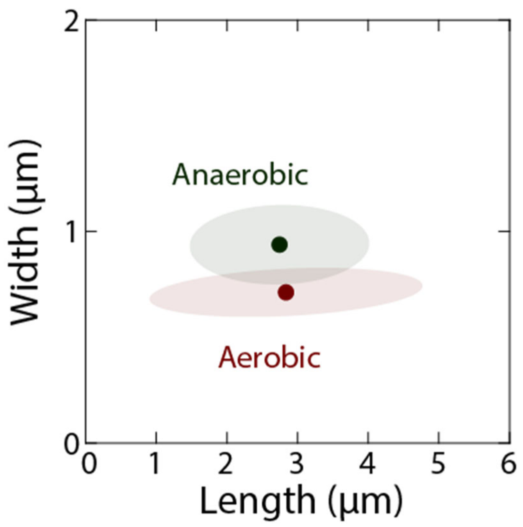


**Figure S9: Cells sampled from the edge of anaerobically grown colonies are wider and shorter than in aerobically grown colonies.** *n*>50 cells were quantified for each condition.

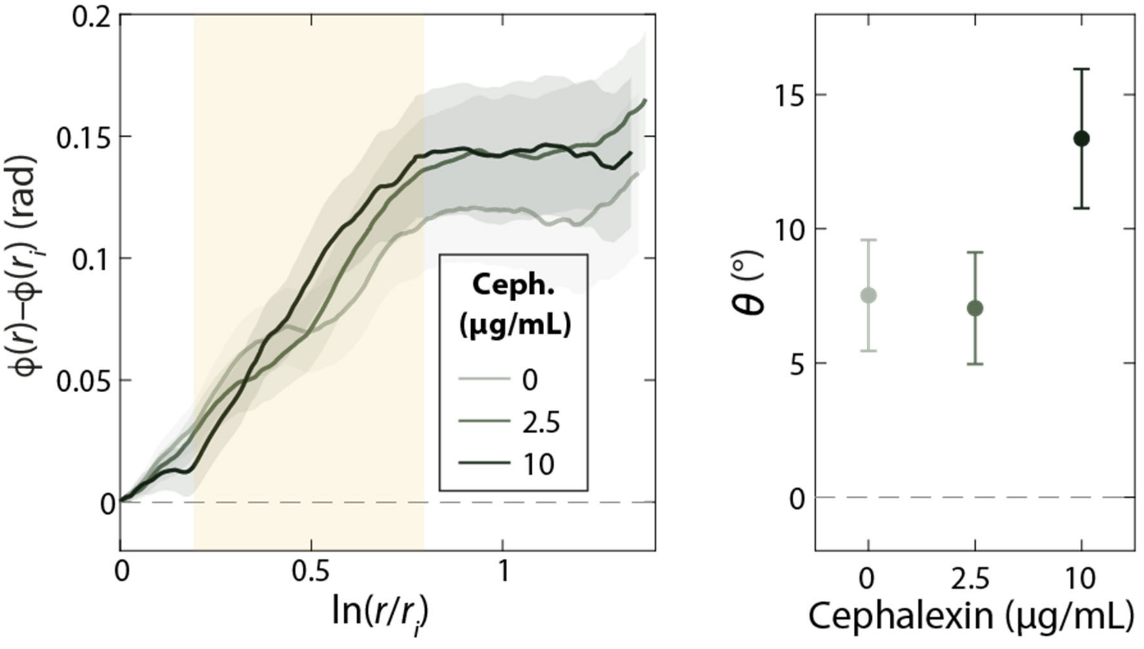


**Figure S10: Cephalexin treatment increases colony chirality.**

Shown is a biological replicate of the experiment in Fig. 7A. Left: mean rotation of sector boundaries of aerobically grown DH5α wild-type colonies with various concentrations of cephalexin. Right: the chiral angle increases with increasing cephalexin concentration. Each data point is the average of $n \geq 5$ colonies. Error bars represent 1 standard deviation.
